# Nicotinamide mononucleotide redox cofactor system enables aldehyde accumulation in *Escherichia coli*

**DOI:** 10.1101/2020.05.11.089136

**Authors:** Kelly N. Richardson, William B. Black, Han Li

## Abstract

It is challenging to biosynthesize industrially important aldehydes, which are readily consumed by the numerous alcohol dehydrogenases (ADHs) in cells. In this work, we demonstrate that a nicotinamide mononucleotide (NMN^+^)-dependent redox cofactor cycling system enables aldehyde accumulation in *Escherichia coli* crude lysates and whole cells. By specifically delivering reducing power to a recombinant enoate reductase, but not to endogenous ADHs, we convert citral to citronellal with minimal byproduct formation (98% and 83% product purity in crude lysate- and whole cell-based biotransformation, respectively). We envision the system’s universal application to lower the noise in biomanufacturing by silencing the host’s metabolic background.

## Introduction

Controlling the flow of mass and energy through a cell’s metabolism is a central theme of biocatalyst development. Limiting side reactions is key to developing processes that are efficient and economically viable in the increasingly petroleum-dependent world^1,2^. The need for controlling side reactions is particularly evident in the biorenewable production of aldehydes, which are used industrially for flavors, odorants, and as a precursor to plastics and resins^3-5^. Engineering microorganisms to accumulate aldehydes has proven difficult due to the high activity and broad substrate ranges of a myriad of NAD(P)H-dependent alcohol dehydrogenases (ADHs) in the cell, which readily reduce aldehydes to alcohols^6-9^.

To overcome this challenge, various techniques have been employed. In *Saccharomyces cerevisiae*, pathway modules have been compartmentalized in organelles to physically isolate the accumulating aldehydes from competing enzymes^10,11^. In *Escherichia coli*, screening, characterization, and disruption of aldehyde reductases and ADHs has enabled the engineering of several aldehyde accumulating strains^3,12^. For example, Rodriguez and coworkers generated an aliphatic aldehyde accumulating strain by screening 44 candidate aldehyde reductases and ultimately identifying and disrupting 13 genes^3^. Similarly, Kunjapur and coworkers developed an aromatic aldehyde accumulating *E. coli* strain, termed reduced aromatic <u>a</u>ldehyde <u>re</u>duction (RARE) strain, by disrupting six aldehyde reductases^12^. They then used the RARE strain in the production of benzaldehyde and vanillin^12^. While both aldehyde accumulating strains enabled significantly increased aldehyde titer and yields, success of these strains is substrate-dependent^13,14^, and both groups have recognized that uncharacterized consumption of aldehydes still persist^3,12^. Furthermore, the process of identifying, characterizing, and screening the numerous reductases is time consuming and labor intensive, a task which becomes even more arduous when engineering non-model organisms without well understood metabolisms.

Recently, we demonstrated the use of a nicotinamide mononucleotide (NMN^+^)-dependent orthogonal redox cofactor system to control pathway flux without the need to genetically disrupt competing pathways^15^. Here, we demonstrate that this system can serve as a new tool for aldehyde production by curtailing electron supply to ADHs. This concept was previously explored by Paul and coworkers where they employed chemically reduced biomimetic cofactor to decrease ADH activity during *in vitro* production of citronellal^16^, a terpenoid aldehyde with wide usage in the flavor and fragrance industries, as the precursor of nonracemic menthol, and as an insect repellant^17,18^. However, because this system did not utilize a method of cofactor regeneration, an equimolar supply of chemically reduced cofactor was required. Moreover, its application to *in vivo* biosynthesis was not achieved.

In this work, we first demonstrate that the NMN^+^-based orthogonal redox cofactor system can enable citronellal production in crude lysate-based, cell-free biotransformation without the need to identify or disrupt competing ADHs in *E. coli*. Next, we used the modular crude lysate system to rapidly prototype and optimize the process, resulting in a productivity of 76 mg/L citronellal in 1 hour, with a product purity to 98%. Finally, the optimized crude lysate system was used to inform whole-cell biotransformation in *E. coli*, which produced 33 mg/L citronellal with 83% product purity. In contrast, biotransformation processes relying on the natural cofactors nicotinamide adenine dinucleotide (phosphate) (NAD(P)^+^) only yielded the alcohol byproducts, namely citronellol, nerol, and geraniol. We envision this approach may be adapted for different target products and in various organisms with relative ease, since the successful application of the orthogonal redox cofactor technology does not depend on comprehensive knowledge of the host’s endogenous metabolic activity.

## Results and Discussion

### Design of the NMN^+^-dependent citronellal production scheme

We employed *Pseudomonas putida* enoate reductase XenA to reduce the activated C=C double bond in citral to yield citronellal (Figure 1). The reducing power required by XenA is provided by glucose oxidation, which is catalyzed by *Bacillus subtilis* glucose dehydrogenase (GDH). Depending on whether the wild type GDH (GDH WT) or engineered GDH (GDH Ortho) is used, NAD(P)^+^ or the noncanonical cofactor NMN^+^ can be reduced. While XenA can accept reducing power in the forms of both NADP(H) and reduced NMN^+^ (NMNH), endogenous ADHs only accept NAD(P)H. As a result, when GDH WT is used, all reactions catalyzed by XenA and ADHs can occur, resulting in the production of not only citronellal, but also alcohol byproducts, namely geraniol, nerol, and citronellol (Figure 1A). On the other hand, we hypothesized that when GDH Ortho is used, NMN^+^ will be the only cycling cofactor which delivers electrons specifically to XenA, yielding citronellal as the predominant product (Figure 1B).

**Figure 1:**
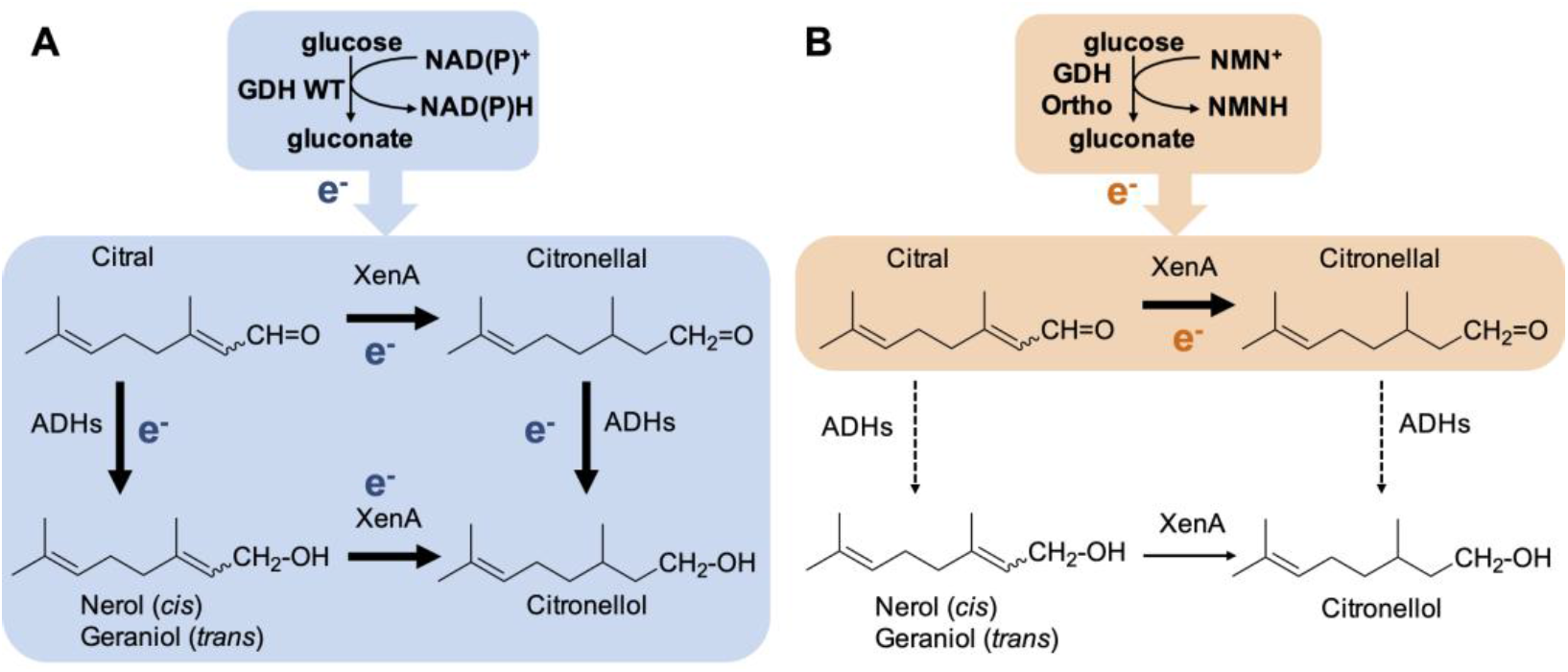
Reaction scheme for citronellal production. A two-part redox cofactor cycling system facilitates the conversion of citral to citronellal through the action of *Pseudomonas putida* XenA. Redox cofactors are continuously regenerated by the glucose dehydrogenase, GDH, from *Bacillus subtilis*. (A) When the wild type GDH is supplied, the native cofactors nicotinamide dinucleotide (phosphate), NAD(P)^+^, are reduced. This supplies the reducing power to both XenA and *E. coli*’s endogenous alcohol dehydrogenases (ADHs), resulting in the formation of alcohols (nerol, geraniol, citronellol) as the final products. (B) When a nicotinamide mononucleotide (NMN^+^)-specific GDH, GDH Ortho, is introduced, reducing power is only delivered to XenA, which can readily receive NMNH. Endogenous ADH activity is silenced since ADHs cannot receive NMNH. Therefore, citronellal, the target product, will accumulate.

### Citronellal production in crude lysate-based, cell-free biotransformation

We first tested our hypothesis in crude lysate-based biotransformation. Crude lysate-based biotransformation serves as a great tool in biocatalysis because it has a cell-like construction and metabolism, but it is freed from the liability of maintaining living cells^19-21^. However, just like in whole cells, controlling pathway flux and limiting non-specific reactions is critical for efficient product formation in crude lysate-based biosynthesis^16^.

The orthogonal redox cofactor system was built in a modular fashion. *E. coli* crude lysates containing overexpressed different variants of GDH (on plasmids pEK101 or pLZ216, Table S1) or XenA (on plasmid pEK102) were generated separately and subsequently mixed-and-matched to yield a complete cycling system for each experimental condition. The *E. coli* host, MX102 (Table S1), also contains knockouts to block reducing power generation from glucose by the host’s native metabolism^15^. When XenA and GDH WT lysates were mixed at a 1:1 ratio, and with the supplementation of no additional cofactor, 1 mM NAD^+^, or 1 mM NADP^+^, citronellal only transiently accumulated to a low level (<10 mg/L) before being rapidly consumed (Figure 2A). Notably, the substrate citral also disappeared rapidly (Figure S1). These results highlight the challenge in citral biotransformation into citronellal since both the substrate and the product are aldehydes.

**Figure 2:**
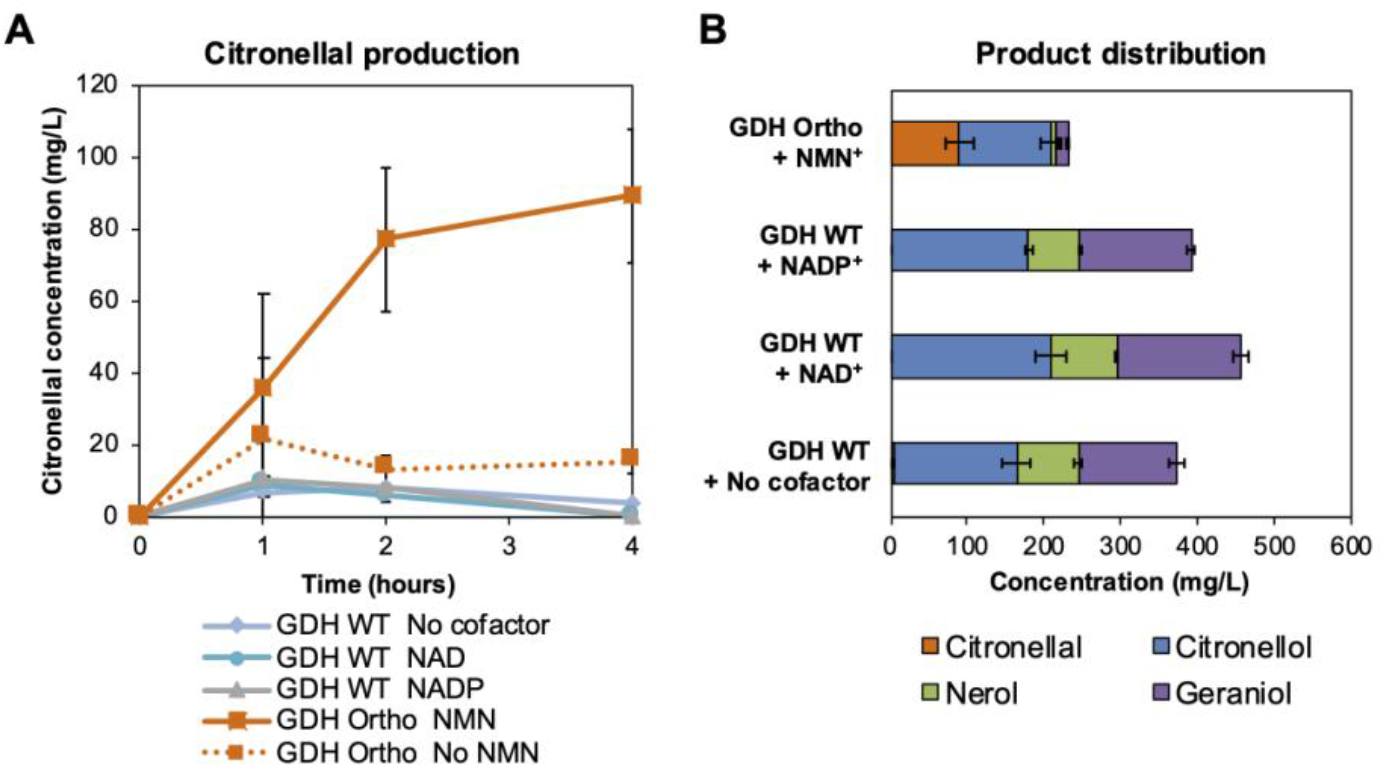
NMN^+^-dependent orthogonal redox cofactor system enables citronellal accumulation. Crude *Escherichia coli* lysates enriched with XenA and wild type *Bacillus subtilis* GDH (GDH WT) or engineered GDH (GDH Ortho) was used to convert citral to citronellal. (A) When GDH WT was used with XenA, citronellal only transiently accumulated to low levels. When GDH Ortho was used with XenA, citronellal was able to accumulate. (B) Product distribution of cycling reactions. The reactions contain 4.5 mg/L lysates, 12 mM magnesium acetate, 10 mM ammonium acetate, 130 mM potassium acetate, 10 mM potassium phosphate, 200 mM D-glucose, 1 mM of oxidized cofactor, and 500 mg/L citral. Reactions were incubated at 37 °C. Values are an average of at least three replicates, and the error bars represent one standard deviation.

Importantly, when GDH Ortho was used in place of GDH WT, 89 mg/L of citronellal was produced with a 39% product purity (the percentage of citronellal in the sum of all products including citronellal, geraniol, nerol, and citronellol) when 1 mM of NMN^+^ was supplemented (Figure 2A, B). When NMN^+^ was omitted, citronellal production was not significant, demonstrating that aldehyde accumulation was indeed NMN^+^-dependent (Figure 2A). Remarkably, this was achieved without disrupting any aldehyde reductases or ADHs genes.

### Optimizing the crude lysate-based biotransformation

Although the crude-lysate system was able to accumulate citronellal, approximately 60% of the products were still alcohols, with the major byproduct being citronellol (119 mg/L) (Figure 2E). We hypothesized that although GDH Ortho specifically generates NMNH from glucose, NAD(P)H is still generated from other substrates present in crude lysates, such as lipids, amino acids, and pyruvate, and that this generation of NAD(P)H was enabling alcohol formation. Although the amount of NAD(P)H generated through these background activities may be small, the highly active, endogenous ADHs are able to use it efficiently. Instead of screening and disrupting the numerous pathways in *E. coli* that can generate NAD(P)H, we sought to increase XenA activity to outcompete ADHs, and to optimize the balance between GDH Ortho and XenA.

We capitalized on the modularity of the system to rapidly prototype various ratios of XenA:GDH Ortho lysates^22,23^, with the total amount of lysates held constant. The 1:1 ratio tested in the initial proof-of-concept (Figure 2) only produced 24 mg/L of citronellal with a product purity of 44% in 1 hour (Figure 3A). While increasing the proportion of GDH Ortho had a detrimental effect, shifting the ratio towards XenA resulted in increased citronellal production; this is consistent with our hypothesis that the productivity was impeded by poor relative XenA activity (Figure 3A). The 7:1 (XenA:GDH Ortho) ratio produced 69 mg/L of citronellal and a 78% product purity, with 16 mg/L citronellol (Figure 3A) and undetectable levels of geraniol and nerol (data not shown) produced. Increasing the XenA:GDH Ortho ratio to 11:1 further diminished citronellol production and increased product purity of citronellal to 86% (Figure 3A). However, this increase in purity came at the loss of citronellal production to 65 mg/L (Figure 3A), likely because GDH Ortho became limiting in this condition. These results indicated that the cycling system functioned most effectively at a 7:1 XenA:GDH Ortho ratio. When we added an ultracentrifugation step in lysate preparation to further remove interfering enzymes which are associated with the membranes or present in high molecular-weight protein complexes, the productivity and purity was further improved (Figure 3B). When ultracentrifuge lysates were tested at the 7:1 XenA:GDH Ortho ratio, 75 mg/L citronellal was produced in 1 hour with a 98% product purity. Alcohol byproduct formation was blocked, with < 2 mg/L citronellol and undetectable levels of geraniol and nerol produced (Figure 3B).

**Figure 3:**
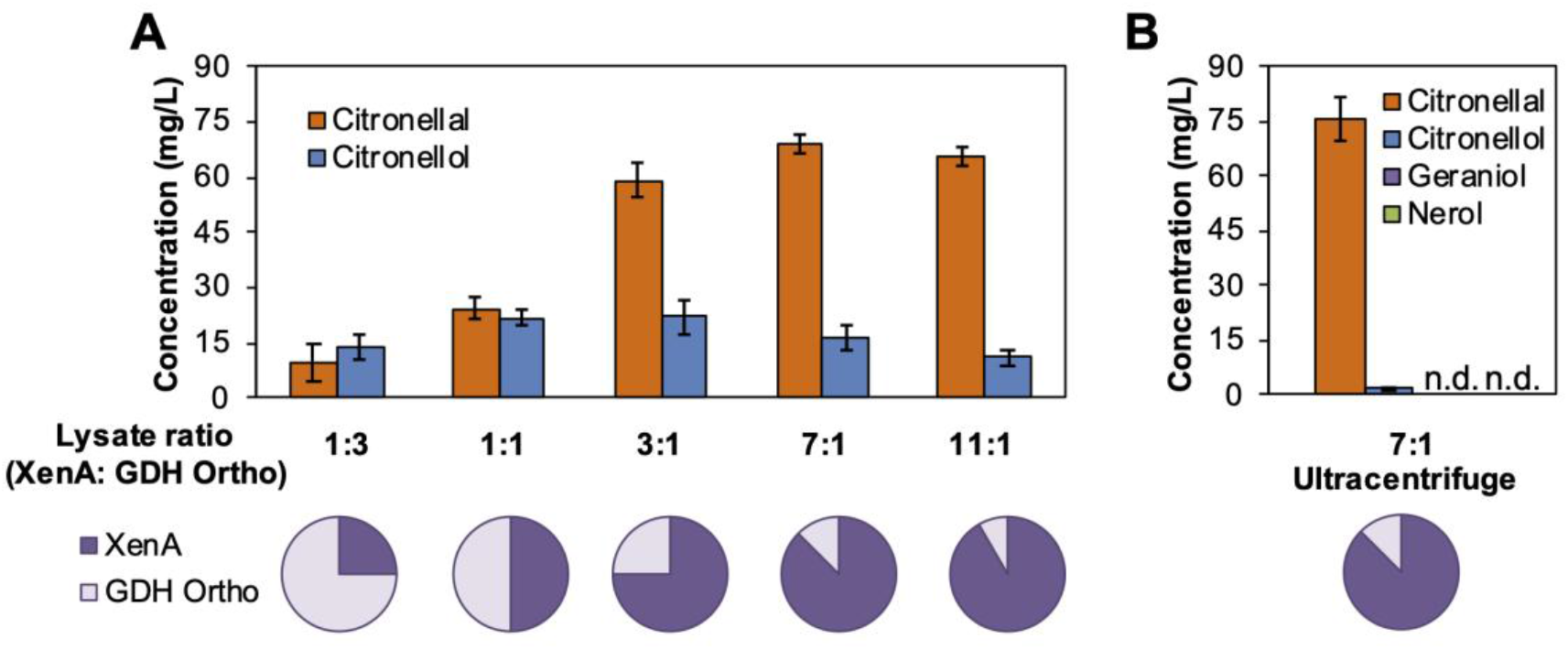
Optimizing crude lysate-based citronellal production. (A) Increasing the relative amount of XenA- to GDH Ortho-enriched *E. coli* crude lysate enabled increased citronellal production and specificity. (B) Further clarifying the lysate with an ultracentrifuge further improved the performance, yielding a >98% pure production of citronellal. The reactions contain 4.5 mg/L lysates, 200 mM D-glucose, 1 mM of oxidized cofactor, and 500 mg/L citral and were incubated at 37 °C for 1 hour. Values are an average of at least three replicates, and the error bars represent one standard deviation. n.d., not detected.

### Citronellal production in *E. coli* whole cells

We next examined the ability of the orthogonal cofactor system to produce citronellal *in vivo*. Consistent with the cell-free process, when resting *E. coli* whole cells were supplied with XenA (on plasmid pEK101, Table S1), GDH WT (on plasmid pSM107), and a glucose facilitator, *Zymomonas mobilis* Glf (on plasmid pSM109) which transports glucose into the cell, endogenous ADH activity completely consumed citral and citronellal, resulting in the production of only geraniol, nerol, and citronellol (Figure 4A). Importantly, when GDH Ortho was used to replace GDH WT, aldehyde accumulation was achieved (Figure 4B).

**Figure 4:**
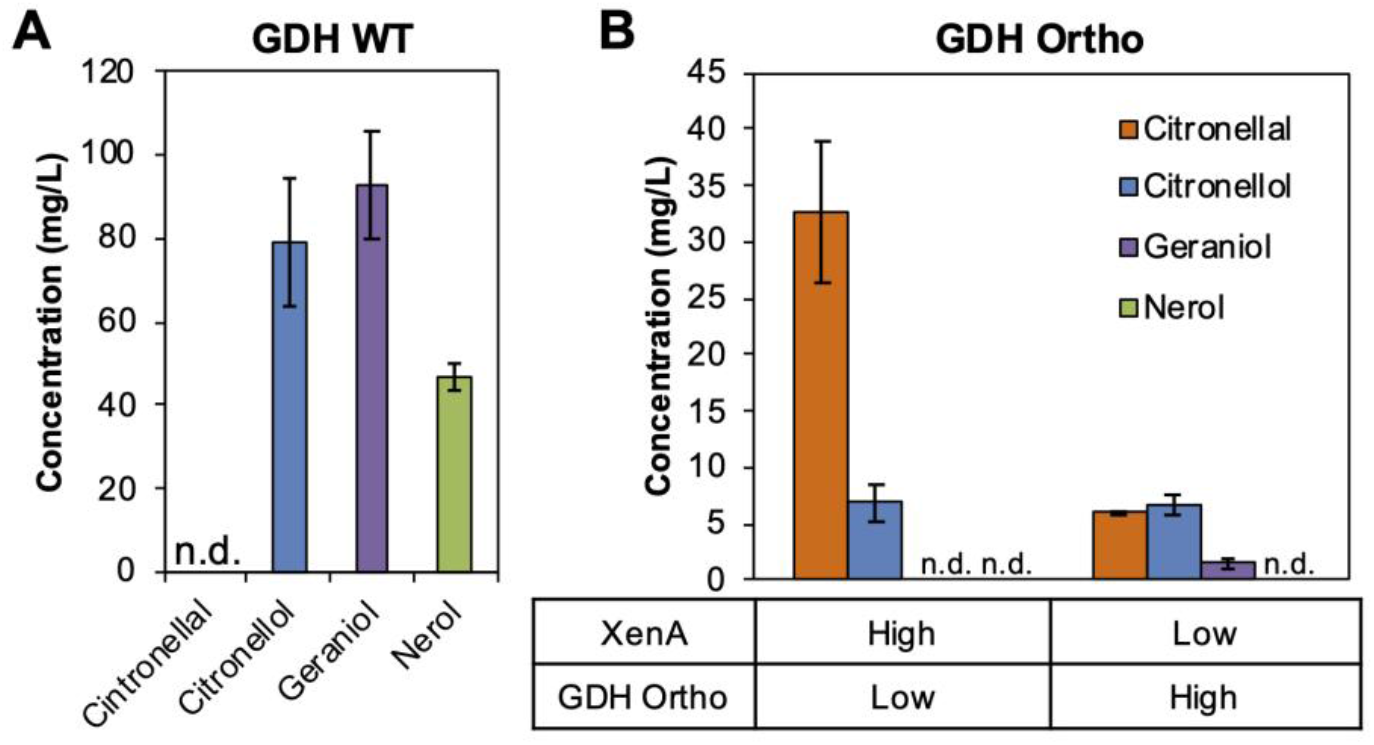
Orthogonal cofactor system enables specific aldehyde accumulation in *E. coli* whole cells. A cofactor cycling system which cycles NAD(P)^+^ (*via* GDH WT) or NMN^+^ (*via* GDH Ortho) was introduced to resting *E. coli* cells to enable the conversion of citral to citronellal by XenA. (A) With GDH WT, only alcohol byproducts derived from citral and citronellal were produced. (B) With GDH Ortho, citronellal accumulation was achieved. High XenA expression relatively to GDH Ortho resulted in higher citronellal productivity and product purity. Whole-cell biotransformation was performed with resting *E. coli* cells at an OD600 of 10 with 200 mM D-glucose, 2 mM NMN^+^, and 500 mg/L citral, at 30 °C for 3 hours while shaking at 250 r.p.m. Values are an average of at least three replicates, and the error bars represent one standard deviation. n.d., not detected.

Previously, crude-lysate based biotransformation has been shown as a powerful tool to prototype and inform pathway design in whole-cells^24,25^. Here, we demonstrated the benefit of maintaining a higher XenA:GDH Ortho ratio is translatable from *in vitro* to *in vivo*. We introduced the orthogonal cofactor cycling system using two different configurations (Figure 4B). When high XenA expression was paired with low GDH Ortho expression (on plasmids pEK101 and pSM106, respectively. Figure S2, Table S1), 33 mg/L of citronellal was produced with a product purity of 83% (Figure 4B). Conversely, pairing high GDH Ortho expression with low XenA expression (on plasmids pLZ216 and pLZ217, respectively) caused a significant decrease in citronellal production (6 mg/L), and a poor product purity of 42%. In both conditions, the major byproduct citronellol was formed at 7 mg/L.

The orthogonal redox cofactor system is highly complementary to the commonly used host engineering methods. For example, while the orthogonal redox cofactor system bypassed the laborious process of knocking out the numerous, highly promiscuous ADHs and aldehyde reductases^3,12^, efforts to genetically eliminate the fewer, more substrate specific, and more easily identifiable competing enzymes that consume citral and citronellal (namely aldehyde dehydrogenases and citral lyase) may further improve biotransformation performance^17,26^.

## Conclusion

This work demonstrates the use of an NMN^+^-dependent orthogonal redox cofactor system as a facile method to develop biocatalysts for aldehyde production. As a proof-of-concept, we achieved effective conversion of citral to citronellal (both are aldehydes) by silencing the highly active ADHs in both *E. coli* crude lysates and whole cells. Notably, we decreased the level of alcohol byproducts in crude lysate-based biotransformation from 60% to 2%, by only optimizing the components in the orthogonal redox loop, and without relying on any information about the host’s natural metabolism. The ease in optimization highlights the advantage of an insulated, orthogonal metabolic system. Compared to methods which aim to achieve insulation through physical compartmentalization^10,11^, the orthogonal redox cofactor technology chemically insulates aldehydes from competing ADHs. This avoids the need to target and transport enzymes and chemicals into natural or artificial organelles, and is inherently beneficial in crude-lysate based biotransformation where cells are homogenized and the spatial organization may be lost. As more enzymes are engineered to efficiently utilize NMN(H), the methods established here may be readily adapted to produce various chemicals and in different microbial chassis.

## SUPPORTING INFORMATION

Experimental methods, plasmids and strains used in this study (Table S1), accession numbers for proteins used in this study (Table S2), citral consumption in crude lysate-based biotransformation (Figure S1), and expression levels of different vectors as measured by enzyme activity in crude lysate (Figure S2).

## AUTHOR CONTRIBUTION

H.L. conceived the work. K.N.R. and W.B.B. performed the experiments. H.L., K.N.R., and W.B.B. analyzed the data. H.L., K.N.R., and W.B.B. wrote the manuscript.

## NOTES

The authors declare no competing financial interest.

## ACKNOWLEDGMENT

H.L. acknowledges support from University of California, Irvine, the National Science Foundation (NSF) (award no. 1847705), and the National Institutes of Health (NIH) (award no. DP2 GM137427). W.B.B. acknowledges support from Graduate Assistance in Areas of National Need fellowship funded by the U.S. Department of Education. The content is solely the responsibility of the authors and does not necessarily represent the official views of the National Institutes of Health or the NSF.

## Notes

### Competing Interest Statement

The authors have declared no competing interest.

