## Supplementary information for "Nicotinamide mononucleotide redox cofactor system enables aldehyde accumulation in *Escherichia coli*"

### Supporting Information

**A. Methods**

**B. Supporting Figures and Tables**

**Figure S1:** Citral consumption in crude lysate-based biotransformation

**Figure S2:** Expression levels of different vectors as measured by enzyme activity in crude lysate

**Table S1:** Plasmids and strains used in this study

**Table S2:** Accession numbers for proteins used in this study

**C. References**

### A. Methods

#### Plasmids and Strains

All plasmids and strain MX102 were taken from previous work in our lab [1] (Table S1). All plasmids were transformed by electroporation. When multiple plasmids were used in a strain, they were transformed simultaneously. When applicable, the following antibiotic concentrations were used, unless stated otherwise: 100 mg/L ampicillin, 50 mg/L kanamycin, 50 mg spectinomycin, or 20 mg/L chloramphenicol.

#### Crude Lysate Biotransformations

Cell-free reactions were performed with crude *E. coli* lysates in a modular system. In this system, each protein was overexpressed and processed separately. Then, different lysates were mixed at controlled ratios to complete the full system. *E. coli* strain MX102 was transformed with a plasmid expressing XenA (pEK102), GDH WT (EK101), or GDH Ortho (pLZ216). 4 mL seed cultures containing 2xYT medium supplemented with 1% w/v mannitol and appropriate antibiotics were incubated at 30 ℃ while shaking at 250 r.p.m. for 16 hours. Next, 0.5% v/v seed cultures were used to inoculate 250 mL of 2xYT medium supplemented with 1% w/v mannitol and 200 mg/L ampicillin in a 500 mL baffled shake flask. Cultures were incubated at 30 ℃ while shaking at 250 r. p. m. for 4 hours. Protein expression was induced with 0.5 mM IPTG and 0.1% w/v arabinose. Cultures shook for an additional 10 hours under the same conditions. To harvest and process the cells, cultures were centrifuged at 4000 RCF for 20 minutes at 4 ℃. The resulting cell pellets were washed 3 times in 250 mL of ice-cold wash buffer containing 120 mM potassium acetate, 28 mM magnesium acetate, and 20 mM tris base pH 8.2. After washing, cells were resuspended in 0.7 mL of wash buffer per 1 g wet-cell-weight of pelleted cells. The resuspended cells were then lysed by French press. The cell lysate was clarified by centrifugation twice at 20,000 RCF for 20 minutes at 4 ℃. When stated in the text, an additional ultracentrifugation step was applied by centrifuging the supernatant from the preceding centrifugation step at 30,000 RCF for 30 minutes at 4 ℃. The clarified lysate was aliquoted into 1.5 mL microcentrifuge tubes and stored at -80 ℃ until use. The total protein concentration in the lysate was quantified by Bradford Assay.

Cell-free reactions were performed at a 1mL working volume in 2 mL glass vials sealed with a PTFE-lined cap at 37 ℃ without shaking. Clarified cell lysate was thawed on ice. The XenA-enriched lysate was mixed with the GDH WT- or GDH Ortho-enriched lysate at various ratios to create a complete cycling system. The total lysate protein concentration was maintained constant at 4.5 mg/L in the final reaction mixture. Lysate mixture was spiked into pre-warmed, concentrated assay buffer to start the reaction. The working concentration and composition of the reaction mixture was 12 mM magnesium acetate, 10 mM ammonium acetate, 130 mM potassium acetate, 10 mM potassium phosphate, 200 mM D-glucose, 1 mM of oxidized cofactor, and 500 mg/L citral. Samples were taken intermittently for analysis. Samples were extracted with an equal volume of chloroform containing 200 mg/L octanol as an internal standard. The organic fraction was used for GC-FID analysis, as detailed below.

#### Resting Whole-Cell Biotransformation

Three plasmids, one expressing XenA (selected from pEK102 or pLZ217), one expressing one of the GDH variants (selected from pLZ216, pSM106, or pSM107), and pSM109 expressing a glucose transport facilitator were transformed into MX102. 4 mL seed cultures containing 2xYT medium supplemented with 1% w/v mannitol and appropriate antibiotics were incubated at 30 ℃ while shaking at 250 r.p.m. for 16 hours. Next, 0.5% v/v seed cultures were used to inoculate 100 mL of 2xYT medium supplemented with 1% w/v mannitol in a 250 mL baffled shake flask. Cultures were incubated at 30 ℃

while shaking at 250 r. p. m. for 4 hours. Protein expression was induced with 0.5 mM IPTG and 0.1% w/v arabinose. Cultures were shaken for an additional 10 hours under the same conditions. Cells were collected by centrifugation in 50 mL conical tubes for 15 minutes at 20 ℃ at 3,750 r.p.m.. The supernatant was discarded, and the cells were washed three times with 50 mL of room temperature 100 mM potassium phosphate (pH 7.4), followed by resuspension in room temperature 100 mM potassium phosphate at pH 7.4 to a standardized $OD_{600}$ = 50. 1 mL of washed cells were spiked into 4 mL of concentrated assay buffer in a 15 mL conical tube. The working concentration and composition of buffer contained 100 mM potassium phosphate (pH 7.4), 200 mM D-glucose, 2 mM $NMN^+$, and 0.5 g/L citral. The tubes were sealed tight and mounted horizontally in an incubating shaker. The tubes were incubated at 30 ℃ while shaking at 250 r.p.m. for 3 hours. After incubation, 0.5 mL of cell suspension was pelleted, and 200 µL of supernatant was extracted with an equal volume of chloroform containing 200 mg/L octanol as an internal standard. The organic fraction was analyzed by GC-FID, as detailed below.

**GC-FID Analysis**

All GC analysis was performed on an Agilent 6850 (Agilent Technologies) equipped with an FID. An Agilent DB-WAXetr capillary column (30 m x 0.56 mm x 1 µm) was used for separation. The inlet and detector were held at 250 and 260 ℃, respectively. The GC was operated in constant flow mode with a flowrate of 3 mL/min. Helium was used as the carrier gas. Air and hydrogen were supplied to the FID at 350 and 40 mL/min, respectively. All gasses were purchased from Airgas. 5 µL of sample was injected with a 2:1 split ratio. The oven was held at 150 ℃ for 15 minutes, then ramped to 240 ℃ at a rate of 20 ℃/minute before equilibration back to 150 ℃. Elution times are as follows: citral, 7.26 and 8.42 minutes; citronellal, 4.08 minutes; citronellol, 8.72 minutes; nerol, 10.04 minutes; geraniol, 11.55 minutes; octanol, 4.59 minutes.

**Determination of Relative Plasmid Expression Levels**

Strain MX102 was transformed with two sets of plasmids. The first set expressed XenA on a $P_{LlacO1}$ ColE1 *ori* vector (pEK102) and GDH Ortho on a $P_{BAD}$ RSF *ori* (pSM106). The second set expressed the same genes, but on the opposite vector (pLZ217 and pEK101 for XenA and GDH Ortho, respectively). Cells were cultured identically as resting whole-cell biotransformations detailed in the previous Methods section. 10 mL of culture was incubated on ice for 15 minutes, and then pelleted by centrifugation at 4 ℃. The supernatant was discarded. The cell pellet was resuspended in 1 mL of ice-cold 35 mM Tris-Cl at pH 8.0. The resuspended cells were lysed by bead beading with glass beads. Lysates were quickly cooled in an ice water bath to remove any heat accumulated from bead beating. The lysate was clarified by centrifugation at 20,000 RCF for 20 minutes at 4 ℃. The clarified lysate was transferred to a fresh 1.5 mL microcentrifuge tube on ice for immediate analysis. The protein concentration in the lysate was determined by Bradford Assay.

To measure the relative expression levels of the vector, GDH Ortho activity was measured by the change in light absorption at 340 nm, due to NMNH formation using a SpectraMax M3 at room temperature. No substrate was supplied to XenA, so change in light absorption is a measure of only GDH Ortho activity, not cycling activity. Lysate was spiked into concentrated assay buffer to start the reaction. The working concentration and composition of assay buffer was 35 mM Tris-Cl at pH 8.0, 140 mM glucose, 2 mM $NMN^+$.

**B. Supporting Figures and Tables**

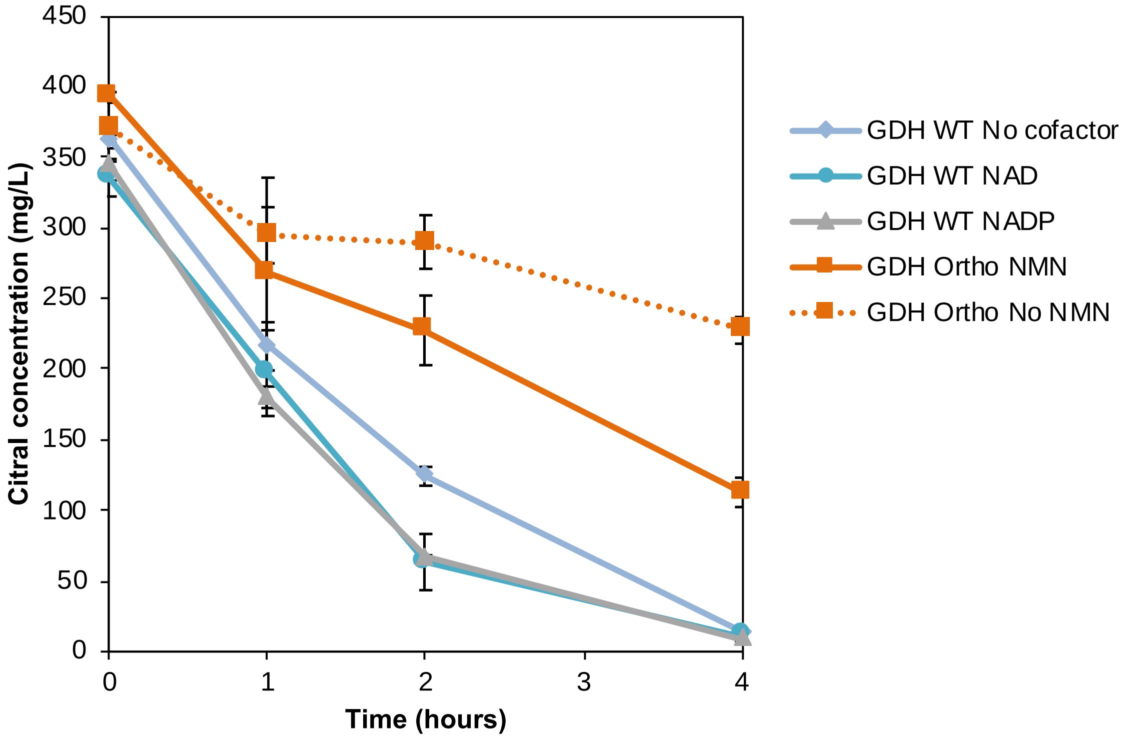


**Figure S1:** Citral consumption in crude lysate-based biotransformation

Crude *Escherichia coli* lysates were used to pair *Pseudomonas putida* enoate reductase XenA with wild type or engineered *Bacillus subtilis* glucose dehydrogenase (GDH WT or GDH Ortho, respectively) to convert citral to citronellal. When GDH WT was supplied, citral was readily consumed by XenA and endogenous aldehyde dehydrogenases. When GDH Ortho was supplied, citral was still present after four hours. However, when GDH Ortho was supplied, citral consumption was not proportional to aldehyde or alcohol product formation (Figure 2B). This indicates aldehyde oxidation and conversion to sulcatone are likely responsible for citral and citronellal degradation, as discussed previously [2]. Confirming this, preliminary results showed the presence of neric acid, geranic acid, citronellic acid, and sulcatone by gas chromatography in the crude lysate reactions (data not shown). Reactions were performed in assay buffer containing 12 mM magnesium acetate, 10 mM ammonium acetate, 130 mM potassium acetate, 10 mM potassium phosphate, 200 mM D-glucose, 1 mM of oxidized cofactor, and 500 mg/L citral. Lysates were mixed at a 1:1 protein ratio. Lysate was spiked into concentrated assay buffer to a final concentration of 4.5 mg/L of protein in solution. Reactions were incubated at 37 °C without shaking. A detailed method can be found in the supplemental information.

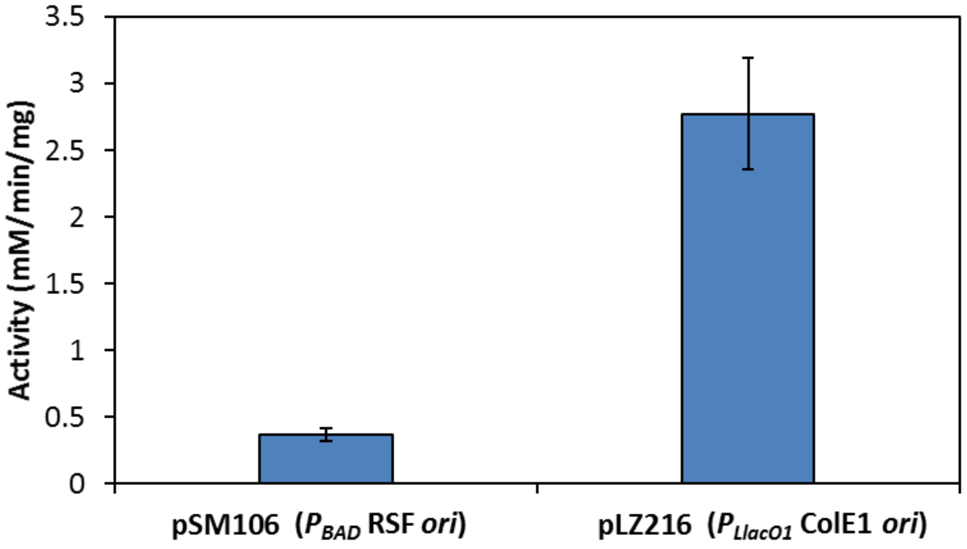


**Figure S2:** Expression levels of different vectors as measured by enzyme activity in crude lysate

To understand the relative expression levels between the $P_{BAD}$ RSF *ori* and $P_{LlacO1}$ ColE1 *ori* plasmids, GDH Ortho's $NMN^+$ reducing activity was measured in crude *Escherichia coli* lysate. The increased $NMN^+$ reducing activity of the $P_{LlacO1}$ ColE1 *ori* vector indicates that the $P_{LlacO1}$ ColE1 *ori* and $P_{BAD}$ RSF *ori* vectors can be treated as a high expression and low expression vectors, respectively, relative to themselves. This methodology was used to vary the expression levels of XenA and GDH Ortho in whole cell biotransformation, as shown in Figure 4. Cells were transformed and cultured identically to the method used for whole cell biotransformation. After cultivating, cells were pelleted, resuspended in ice-cold 35 mM Tris-Cl at pH 8.0, lysed by bead beating, and the lysate was clarified by centrifugation. Lysate was spiked into assay buffer containing 35 mM Tris-Cl, 140 mM D-glucose, and 2 mM $NMN^+$. Activity was determined by change of absorption at 340 nm. A detailed method is discussed in the Methods Section. Values are an average of at least three replicates, and the error bars represent one standard deviation.

**Table S1: Strains and plasmids used in this study**

| **Strains** | **Description** | **Reference** |
|---|---|---|
| MX102 | *E. coli* Δ(*araD-araB*)*567*, Δ*lacZ4787*(::rrnB-3), *λ-*, *rph-1*, Δ(*rhaD-rhaB*)*568*, *hsdR514*, Δ*pncC*, Δ*pgi*, Δ*zwf*, Δ*gntK::kan* | [1] |
| **Plasmids** | **Description** | **Reference** |
| pEK101 | $P_{LlacO1}$*::Bs gdh*, ColE1 *ori*, $Amp^R$ | [1] |
| pEK102 | $P_{LlacO1}$*::Pp xenA*, ColE1 *ori*, $Amp^R$ | [1] |
| pLZ216 | $P_{LlacO1}$*::Bs gdh* I195R-A93K-Y39Q-S17E, ColE1 *ori*, $Amp^R$ | [1] |
| pLZ217 | $P_{BAD}$*::Pp xenA*, RSF *ori*, $Spec^R$ | [1] |
| pSM106 | $P_{BAD}$*::Bs gdh* I195R-A93K-Y39Q-S17E, RSF *ori*, $Spec^R$ | [1] |
| pSM107 | $P_{BAD}$*::Bs gdh*, RSF *ori*, $Spec^R$ | [1] |
| pSM109 | $P_{LlacO1}$*::Zm glf*, p15A *ori*, $Cm^R$ | [1] |

Abbreviations indicate source of genes: *Bs*, *Bacillus subtilis*; *Pp*, *Pseudomonas putida*; *Zm*, *Zymomonas mobilis*

**Table S2:** Accession numbers for proteins used in this study

| Protein Name | Full Name | Protein ID |
|---|---|---|
| *Pp* XenA | NADH: flavin oxidoreductase/NADH oxidase | Q9R9V9 |
| *Bs* Gdh | Glucose 1-dehydrogenase | P12310 |
| *Zm* Glf | Glucose facilitated diffusion porin | P21906 |
